# Inferring Personalized Cell-Cell Communication Networks in Colorectal Cancer with Individualized Causal Discovery

**DOI:** 10.1101/2025.09.20.677543

**Authors:** Aodong Qiu, Binfeng Lu, Gregory F. Cooper, Xinghua Lu, Lujia Chen

## Abstract

Understanding tumor heterogeneity at the resolution of individualized cell–cell communication networks (CCCNs) remains a major computational challenge in precision oncology. Existing inference methods largely rely on population-level correlation and thus fail to capture patient-specific signaling patterns across diverse cell types. To address this limitation, we developed an integrative computational framework combining the nested hierarchical Dirichlet process (nHDP) model for identifying hierarchically structured gene expression modules, with instance-specific Greedy Fast Causal Inference (iGFCI) for inferring individualized CCCNs (iCCCNs) in colorectal cancer (CRC). Applied to large-scale single-cell RNA-seq data from over 625,000 cells, our model successfully decomposed complex gene expression modules GEMs, potentially representing the cell lineage and cellular signaling states, and uncovered iCCCNs across detailed cell subtypes. We further used TCGA bulk RNA-seq data with survival data to validate the clinical relevance of these individualized gene expression module causal interactions, demonstrating their potential as robust prognostic signatures in CRC. Finally, we used principled causal inference methods to search for ligand-receptor pairs that mediate cell-cell communication. This framework enables mechanistic insights into immune evasion. Our computational method represents a significant advance toward realizing personalized oncology, enabling precise patient stratification and identification of actionable biomarkers for improved therapeutic targeting in cancers.

## 1. Introduction

Cell-cell communication (CCC) maintains tissue homeostasis and drives cancer progression through complex signaling networks that vary dramatically between individual tumors^1,2^. Colorectal cancer (CRC) represents the third most common cancer and the second leading cause of cancer death globally, with substantial heterogeneity in tumor composition and functional states that underlie diverse disease progression patterns and treatment responses^3^. Most of CRC tumors do not respond to contemporary immunotherapies, indicating they may exploit immune evasion mechanisms other than ones targeted by current immunotherapies. Understanding previously unknown immune evasion mechanisms by CRC will enhance precision medicine.

Within the tumor microenvironment (TME), diverse cell populations interact through ligand-receptor signaling cascades that reshape cellular states and behaviors, fundamentally influencing immune evasion, metastasis, and therapeutic resistance^4^. However, current approaches for modeling CCCNs predominantly rely on population-level correlational analyses that fail to capture the individualized and causal nature of these interactions. Existing methods such as CellChat^5^, CellPhoneDBv2.0^6^, and Nichenet^7^ focus primarily on cell clusters sharing similar expression patterns and enrichment of ligand-receptor pairs in specific cell clusters as a means to detect potential CCC. However, these methods are unable to explicitly model the downstream impact of transmitted signals or infer causal relationships between cell state changes^8^.

To address these fundamental gaps, we developed a comprehensive Causal Bayesian Network (CBN) learning framework that unveils personalized CCCNs from transcriptomic data (Fig. 1), and we applied the framework on a large collection of CRC tumors to illustrate the framework’s capability of revealing patterns of CCCNs that may underlie heterogeneous TME. We employed nested hierarchical Dirichlet process (nHDP)^9^ model implemented from scGEM package^10^, to deconvolute single-cell transcriptomes from 8 cohorts with over 153 CRC patients and over 625,000 single-cells^11-14^ (Fig. 1A). The nHDP model identifies gene expression modules (GEMs) that capture coordinated transcriptional programs, organizing these GEMs in a hierarchical structure that separates lineage markers from functional cell states (Fig. 1B). Building upon this foundation, we apply individualized causal Bayesian network (iCBN)^15^ inference to construct tumor-specific CCCNs that reveal mechanistic insights into how different cell types causally influence each other within individual tumors (Fig. 1C). Our downstream analysis reveals cross-cell-subtype communication patterns and enables patient stratification into distinct clusters based on their individualized communication network profiles (Fig. 1D). We investigated the molecular mechanisms underlying learned causal edges through ligand-receptor conditional independence testing (Fig. 1E).

**Fig. 1.**
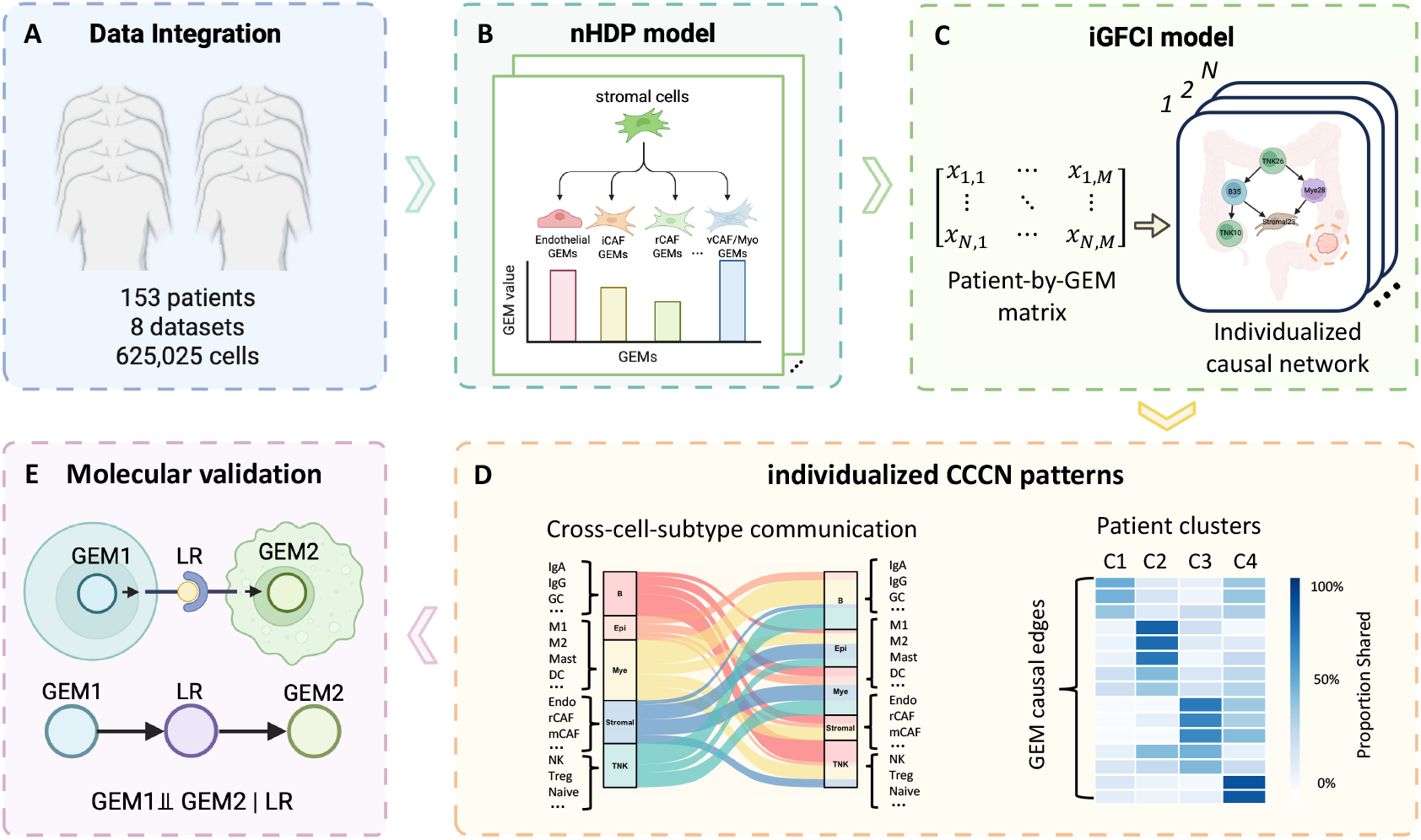
Workflow for this study. **A**. Data integration of single-cell RNA-seq data from 153 patients across 8 datasets, comprising 625,025 cells total. **B**. Application of the nHDP model (implemented in the scGEM package^10^) to identify gene expression modules (GEMs) for each major cell type. Models were constructed for all five cell types: stromal, T and natural killer (TNK) cells, B cells, myeloid cells, and epithelial cells. An example diagram is shown to illustrate subtype-specific GEMs among stromal cells. **C**. Implementation of iGFCI to infer causal relationships between GEMs, generating personalized causal graphs representing CCCNs for each patient. **D**. Analysis of individualized CCCN patterns. A Sankey plot shows cross-cell-subtype communication patterns (left panel). A heatmap shows that CRC patients were clustered (stratification) into four distinct groups based on causal edges from individualized CCCNs (right panel). **E**. Molecular validation of causal edges through searching for ligand-receptor pairs that potentially mediate signal transduction across cells, using the conditional independence test (CIT).

Importantly, this personalized modeling approach enables us to stratify patients within established molecular subtypes into distinct subgroups with different communication patterns and survival outcomes. Our methods discovered fine-grained patient subgroups beyond the conventional patient subtyping of CRC, suggesting that individualized communication networks may serve as novel biomarkers for prognosis and therapeutic targeting. By characterizing the causal architecture of cellular communication at the individual tumor level, our framework represents a novel perspective toward precision oncology and the development of targeted therapeutic strategies.

## 2. Methods

### 2.1. Data integration

We collected and integrated single-cell RNA sequencing data from eight publicly available colorectal cancer cohorts in our previous study^16^, comprising a total of 153 patients and 625,025 high-quality cells. Raw count matrices were preprocessed individually for each dataset using standard quality control (QC) filters, including removal of cells with fewer than 200 expressed genes, more than 20% mitochondrial transcripts, fewer than 500 total UMI counts, or exceptionally high total UMI counts indicative of doublets. All datasets were normalized and log-transformed using Seurat, and batch effects across patients and cohorts were corrected using Harmony^17^. Cells were annotated into five major compartments: TNK cells, myeloid cells, B cells, epithelial cells, and stromal cells, based on canonical marker gene expression. Cells in each major cell type are further annotated into detailed subtypes (Supplementary Figure 1A–D).

### 2.2. Hierarchical GEM discovery via the nHDP model

To identify shared and specialized transcriptional programs within each major cell type, we applied the nHDP model^9^ implemented in the scGEM package^10^. This nonparametric Bayesian framework infers hierarchically structured GEMs. It identifies each cell’s transcriptome as a probabilistic mixture of latent GEMs arranged in a tree-like structure, where deeper nodes represent increasingly specialized transcriptional programs. For each cell *d*, the scGEM model defines a GEM distribution *G*_*d*_ using a recursive stick-breaking construction:

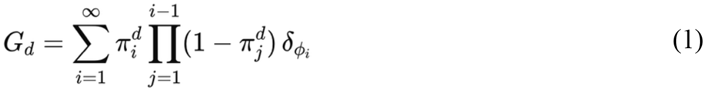

Where 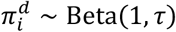 is the stick-breaking proportion assigned to the *i*-th component (GEM) for cell 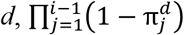 represents the remaining probability mass after allocating to the first *i* − 1 GEMs, 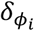 is a Dirac delta function centered at *ϕ*_*i*_, assigning point mass to GEM *ϕ*_*i*_, and *ϕ*_*i*_ ∼ *G* denotes the *i*-th GEM, drawn from a global base distribution *G* over gene expression patterns.

In this study, we preset the tree structure and used a three-level tree structure with five GEMs in the top layer, four children per GEM in the second layer, and three children per GEM in the third layer, yielding a total of *K* = 5+5×4+5×4×3 = 85 GEMs for each major cell type. Under this finite approximation, the GEM distribution becomes:

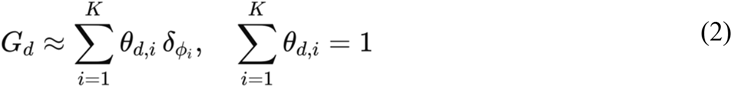

Each GEM *ϕ*_*i*_ is represented by a gene weight vector *β*_*i,g*_. For each cell *d*, we denote the GEM activation weight as a vector *θ*_*d,i*_. We used binarized UMI counts *C*_*d,g*_ ∈ {0, 1} to indicate gene detection. The contribution of cell d to GEM *i* was computed as:

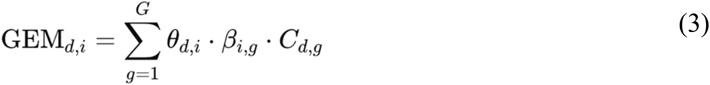

Variational Bayesian inference^18^ with mini-batch updates was used for parameter estimation, and scGEM was applied independently to each major cell type to capture cell-type-specific hierarchical programs.

### 2.3. iGFCI model for searching individualized GEM causal networks

Based on the hypothesis that the expression status of a GEM reflects the state of cellular signal regulating the expression of member genes, we can use GEMs as surrogate variables to represent cellular signaling pathways. Further inference of “causal relationships” between GEMs enables the detection of cell-cell communications. We employed a CBN learning algorithm called instance-specific Greedy Fast Causal Inference (iGFCI)^19^ to learn individualized CCCNs. The input is a patient-by-GEM matrix for each cell type, in which rows represent patients and columns represent GEMs derived from various cell types. This matrix was constructed by aggregating GEM activity across all cells of a given cell type within each patient, summarizing GEM expression at the patient level (aka, pseudo-bulk GEM expression). In order to better capture the functional specificity of each GEM, we annotated the scRNA-seq data from major cell compartments into more refined cellular subtypes (Supplementary Figure 1A–D). We then annotated each GEM based on the dominant expressing cell subtypes, enabling the detection of CCC between highly specialized cell types.

As shown in Fig. 2A, iGFCI operates in two stages^15^. First, population-level model learning: a standard Greedy Fast Causal Inference (GFCI)^20^ algorithm is applied to the full matrix **X** to estimate the population-wide Partial Ancestral Graph (PAG_pop), reflecting the general CCC patterns in CRC patients. Second, for each patient *p*, iGFCI uses the *p*-th row **x**_*p*_ = (*x*_*p*,1_, …, *x*_*p,M*_) to derive an instance-specific Partial Ancestral Graph PAG_IS(*p*) based on PAG_pop and the full matrix **X**. The iGFCI algorithm incorporates context-specific independence (CSI) to prioritize edges that improve model fit specifically for the target instance^15,21^.

**Fig. 2.**
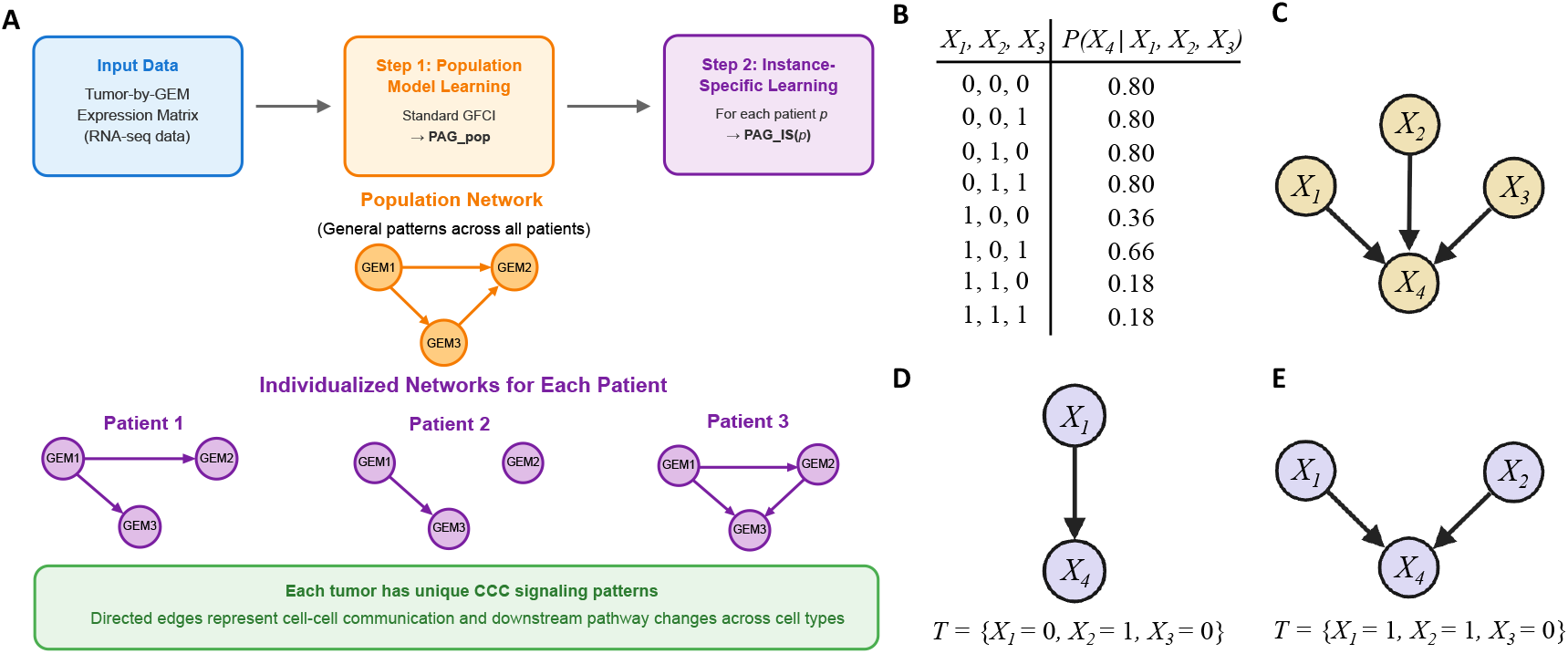
Workflow of iGFCI for learning individualized CCCNs. **A**. The input is a patient-by-GEM expression matrix. Step 1 applies the GFCI algorithm to all patients to learn a population-level PAG_pop, capturing general causal patterns across tumors. Step 2 uses iGFCI to generate instance-specific PAGs, denoted as PAG_IS(*p*), for each patient *p*, revealing individualized causal networks among GEMs. **B**. An example of CSI in CBN. Conditional probabilities of *X*_*4*_ given its parent variables *X*_*1*_, *X*_*2*_, and *X*_*3*_. **C**. CBN learned from the data. **D**. Instance-specific CBN in the context for one instance *T* where *X*_*1*_ = 0, *X*_*2*_ = 1, and *X*_*3*_ = 0. **E**. Instance-specific CBN in the context where *X*_*1*_ = 1, *X*_*2*_ = 1, and *X*_*3*_ = 0.

Fig. 2B-E illustrates an example of CSI. The CBN learned from the full data is shown in Fig. 2C. From the conditional probabilities table (Fig. 2B), we can observe *P*(*X*_*4*_ | *X*_*1*_ = 0, *X*_*2*_, *X*_*3*_) = *P*(*X*_*4*_ | *X*_*1*_ = 0) = 0.80, indicating that *X*_*4*_ ⫫ {*X*_*2*_, *X*_*3*_} | *X*_*1*_ = 0. In other words, when *X*_*1*_ = 0, *X*_*4*_ is conditionally independent of *X*_*2*_ and *X*_*3*_, in a context-specific manner. Therefore, for instance *T* = {*X*_*1*_ = 0, *X*_*2*_ = 1, *X*_*3*_ = 0}, only *X*_*1*_ acts as a direct cause for *X*_*4*_ (Fig. 2D). This can model a biological signaling mechanism where *X*_*1*_ behaves like a gate; when one signal is turned off (e.g., *X*_*1*_ = 0), the other signals (*X*_*2*_, *X*_*3*_) are not able to affect the target (*X*_*4*_). Similarly, *P*(*X*_*4*_ | *X*_*1*_ = 1, *X*_*2*_ = 1, *X*_*3*_) = *P*(*X*_*4*_ | *X*_*1*_ = 1, *X*_*2*_ = 1) = 0.18 (Fig. 2B), so for instance *T* = {*X*_*1*_ = 1, *X*_*2*_ = 1, *X*_*3*_ = 0}, parent nodes of *X*_*4*_ are *X*_*1*_, *X*_*2*_ (Fig. 2E). This demonstrates how iGFCI captures instance-specific causal structure by identifying CSI relationships, which other standard models may miss.

This hybrid constraint-based and instance-specific score-based approach allows iGFCI to identify edges supported in the broader population that also meaningfully explain the data for individual patients. The result is a personalized causal network that reflects unique regulatory and communication relationships among the cells (represented by GEMs) in each patient. To ensure robustness, we performed 100 bootstrap resamplings per patient. Causal edges were retained only if they appeared in more than 20% of bootstrap replicates.

### 2.4. CCCN-based patient stratification and comparison with conventional molecular subtypes

To explore heterogeneity in individualized CCCN, we clustered patients based on their individualized causal graphs (PAG_IS). Specifically, for each patient, we extracted a binary vector encoding the presence or absence of each possible GEM-GEM edge in their PAG_IS, i.e., projecting a patient in the CCCN space. Patients were then grouped into clusters using the K-means clustering algorithm. As a comparison, we also classified patients into the widely used consensus molecular subtype (CMS) subtypes^22^, using the CMScaller^23^ R package applied to patient-level gene expression data. So we can provide a new angle to look into CCC in CRC TME by comparing these two stratification methods.

### 2.5. Survival analysis using TCGA bulk RNA-seq data

To assess the prognostic value of our CCCN-based patient clusters, we extended our analysis to bulk RNA-seq data with associated clinical outcome data from TCGA-COAD and TCGA-READ cohorts. For each GEM, we selected its top 50 contributing genes and used the mean expression level of them to estimate the GEM expression level in bulk tumor samples. For each causal GEM-GEM edge identified in the individualized PAGs, we computed the geometric mean of the two GEM expression scores per patient to quantify the strength of the underlying signaling interaction. Patients were clustered based on these geometric mean scores across causal GEM pairs, and Kaplan–Meier survival curves were generated to evaluate survival differences between clusters.

### 2.6. Investigating molecular mechanisms (ligand-receptor) of causal edges

CCC requires ligand-receptor (LR) pairs to transmit signals in the following fashion: Cell_A_state → LR interaction → Cell_B_state. Here, we use GEM_A from Cell_A and GEM_B from Cell_B as surrogates to represent the state of Cell_A and Cell_B, respectively, leading to GEM_A → LR → GEM_B. This causal chain dictates that when conditioning on LR, GEM_A and GEM_B should be independent. This provides the biological support for inferred causal relationships between GEMs. Following the conventions of the field of using expression values of L and R to model the interaction between a pair of LR^5,6^, we calculated the product [*LR*] ∝ [*L*][*R*] to represent the potential interaction of an LR pair in a TME. We performed conditional independence testing (CIT) using known ligand-receptor (LR) pairs curated from CellPhoneDB v5^24^. For each causal edge between GEM_A and GEM_B, we identified LR pairs where the ligand gene is highly correlated with GEM_A (|Pearson correlation| > 0.3) and the receptor gene is highly correlated with GEM_B (|Pearson correlation| > 0.3). We then applied the Fisher-Z conditional independence test as follows: if the absolute Pearson correlation between GEM_A and GEM_B was greater than 0.3, and the CIT p-value for this GEM pair was greater than 0.05 (null hypothesis is GEM_A ⫫ GEM_B | LR) when conditioning on the expression of the LR pair, we considered this as evidence that the LR pair mediates the dependency between GEM_A and GEM_B.

## 3. Results

### 3.1. The scGEM model reveals hierarchically organized GEM within each cell type

To characterize transcriptomic programs reflecting cellular states in the CRC tumor microenvironment, we applied the nHDP model (scGEM package^10^) to single-cell RNA-seq data across all major cell compartments, including TNK cells, myeloid cells, B cells, epithelial cells, and stromal cells. Here, we use stromal cells as a representative example to illustrate the nested modeling framework and its ability to decompose shared and cell-type-specific gene expression programs. Fig. 3A shows the hierarchical structure inferred by scGEM within the stromal compartment, using a 5-4-3 tree architecture across three layers of increasing specificity. The top layer contains five broad GEMs that capture general transcriptional programs of stromal cells. Each GEM in layer one branches into four child GEMs in layer two, which are further subdivided into three more specific GEMs in layer three, resulting in a total of 85 GEMs for stromal cells. Each node is represented as a pie chart summarizing the average expression of the GEM across annotated stromal subtypes, including endothelial cells, vascular cancer-associated fibroblasts (CAF) with myofibroblastic features (vCAF/Myo), antigen-presenting CAF (apCAF), matrix CAF (mCAF), remodeling CAF (rCAF), and inflammatory CAFs (iCAF).

**Fig. 3.**
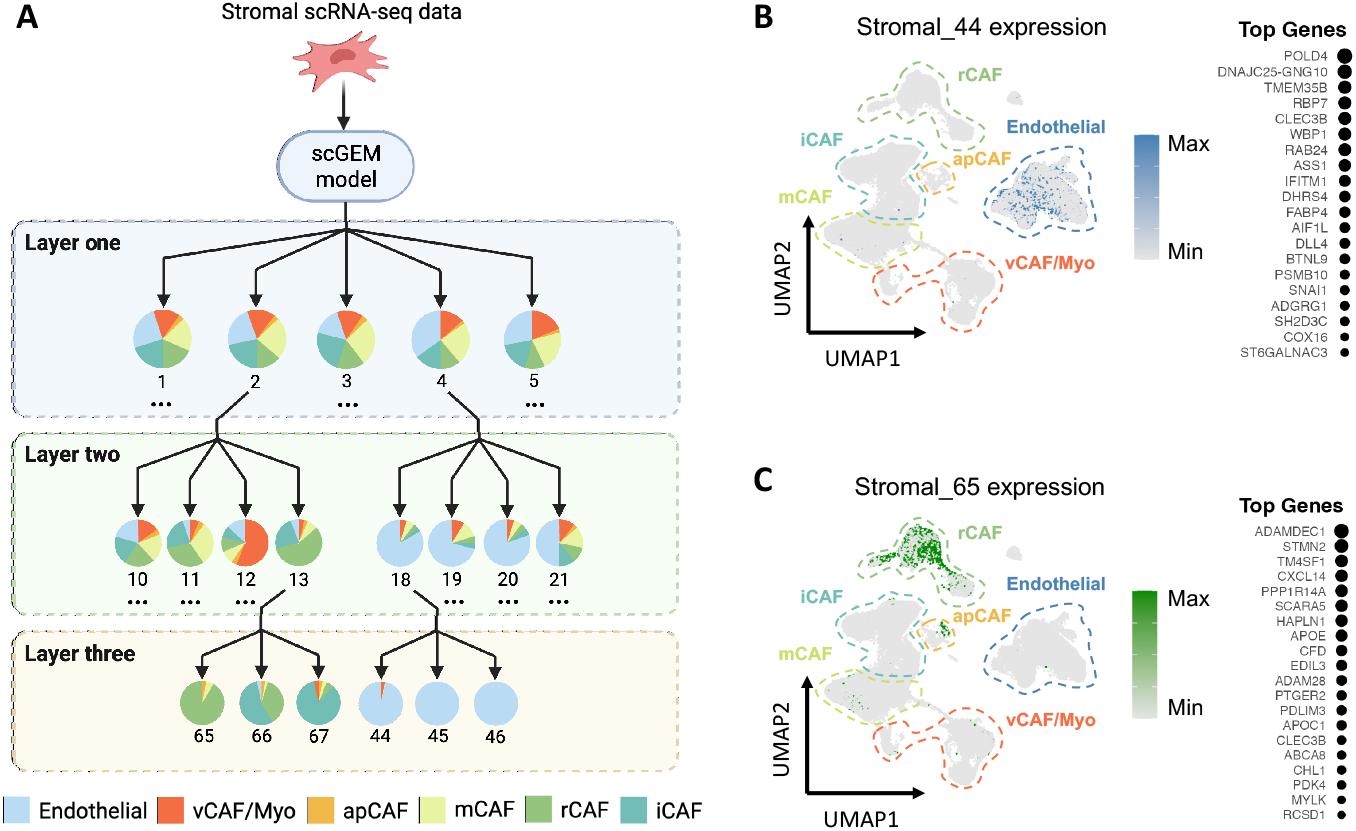
The scGEM model identifies hierarchically structured GEMs in stromal cells. **A**. Schematic representation of the scGEM model structure using stromal cells as an example. The model generates nested tree-structured GEMs across three hierarchical levels (5-4-3 architecture), with separate models constructed for all five major cell compartments (stromal, T/NK, myeloid, B cell, and epithelial cells). Examples shown include pathways from GEM 2 → GEM 13 → GEM 65-67, and GEM 4 → GEM 18 → GEM 44-46, demonstrating how general stromal programs (layer one) give rise to more specialized functional modules (layers two and three). **B**. UMAP visualization of integrated stromal scRNA-seq data showing expression of stromal GEM 44, which is predominantly expressed in endothelial cells. Color intensity represents GEM expression levels, with the top contributing genes for this GEM listed on the right. **C**. UMAP visualization showing stromal GEM 65 expression, which is primarily enriched in rCAF, with the top contributing genes for this GEM listed on the right.

For instance, the stromal_4 → stromal_18 → stromal_44 path reveals endothelial-specific programs. Fig. 3B shows the uniform manifold approximation and projection (UMAP) of stromal cells colored by expression of Stromal_44, which is highly enriched in endothelial cells. This GEM reflects a stressed, angiogenic, and immunomodulatory endothelial state, characterized by genes involved in vascular remodeling (*DLL4, ADGRG1*)^25^, endothelial-to-mesenchymal transition (EndoMT) and invasiveness (*SNAI1*)^26^, and metabolic adaptation and proliferative stress (*FABP4, ASS1, POLD4*)^27-29^. Similarly, the path GEM stromal_2 → stromal_13 → stromal_65 highlights a program that gives rise to a more specialized module stromal_65 enriched in rCAF (Fig. 3C). This GEM captures a distinct remodeling state marked by extracellular matrix remodeling (*ADAM28, EDIL3*)^30,31^, cytokine signaling, and immune modulation (*CXCL14, APOE, PTGER2*)^32-34^.

### 3.2. The iGFCI model reveals patient-specific communication patterns

#### 3.2.1. PAG_pop captures the population-wide communication network

To identify the dominant cell-cell communication pathways shared across CRC patients, we analyzed the PAG_pop from the iGFCI step 1 output, which captures globally consistent causal relationships among GEMs. The PAG_pop revealed frequent signaling pathways spanning multiple compartments (Fig. 4A), including 170 GEM-GEM causal edges (86 cross-cell-type, 84 within cell-type edges). The Sankey plot illustrates these intercellular edges and their respective cell-type origins and targets, highlighting how transcriptomic programs in one cell type potentially regulate those in another, consistent with known cross-cell-type interactions in the TME^4^.

**Fig. 4.**
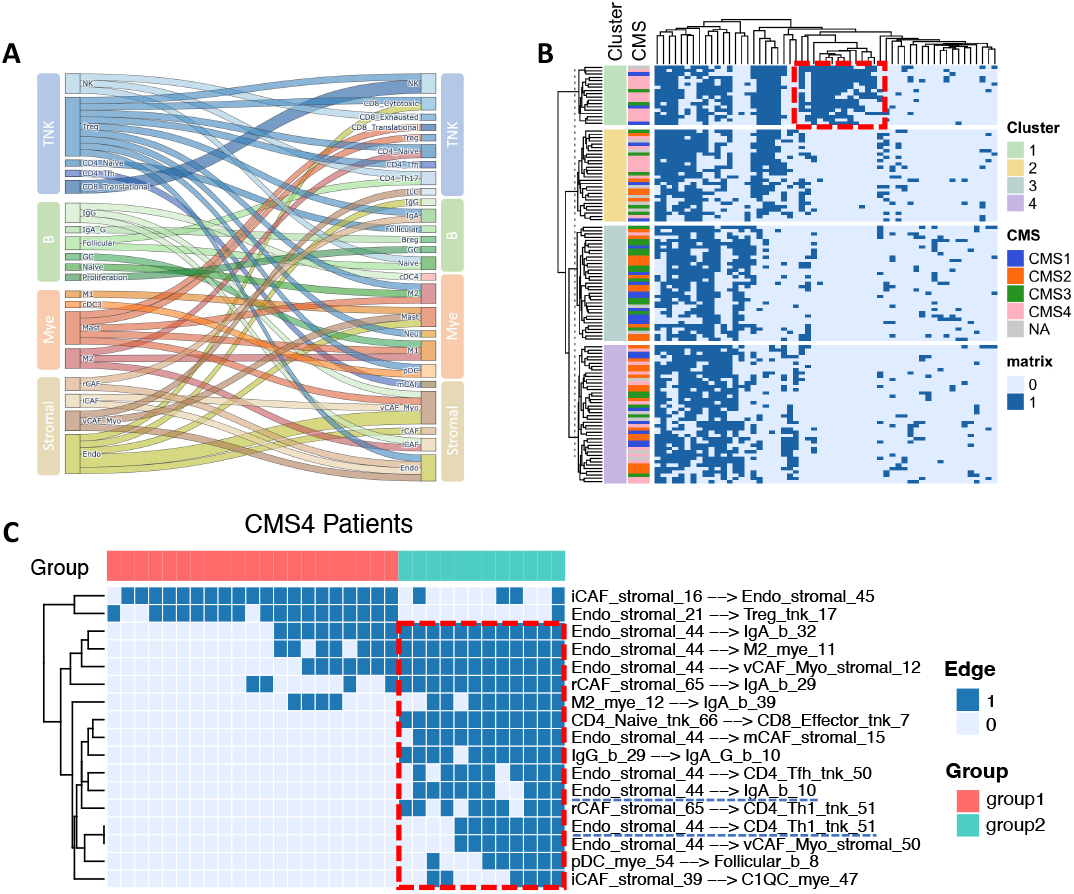
Characterization of GEM Causal Networks Across CRC Patients Using iGFCI. **A**. Sankey plot showing cross-cell-type communication patterns from the population-wide PAG (PAG_pop) inferred by iGFCI Step 1. Each directed edge represents a directed causal edge between GEMs expressed in distinct cell types. **B**. Heatmap of individualized GEM-GEM causal edges from PAG_IS inferred by iGFCI Step 2. Each row corresponds to one patient; each column represents a directed GEM-GEM causal edge. Dark blue entries indicate the presence of a specific edge in a patient’s PAG_IS. Patients were clustered using K-means (K = 4), and annotations include Consensus Molecular Subtypes (CMS1–CMS4) for comparison. **C**. Sub-clustering of CMS4 patients reveals two distinct groups based on their individualized causal edges, demonstrating potential heterogeneity within CMS4 patients.

#### 3.2.2. PAG_IS captures patient-level heterogeneity

While the PAG_pop provides insights into common CCC patterns, it does not account for individual variation. Therefore, applying iGFCI to construct PAG_IS for each patient based on PAG_pop provides insights into studying the individualized immune evasion mechanism. The resulting edge matrices were binary, indicating the presence or absence of each causal edge per patient. Fig. 4B displays the edges in the PAG_IS heatmap across all patients, after filtering out the edges that appear in more than 80% or less than 5% patients. Patients were divided into four clusters using the K-means algorithm, showing distinct causal edge patterns. Notably, our classification appears to be orthogonal to the conventional CMS subtypes, which are mainly based on gene expression profiles. Except for CMS4 samples, which appear to be concentrated in Clusters 1 and 2, other CMS subtypes did not show a strong association with our cluster assignment. This result indicates our method provides a new perspective to stratify CRC tumors, using CCCN patterns instead of the overall gene expression profile.

We further investigate the causal edge patterns in CMS4 samples, which are divided into two clusters Cluster 1 and 2 (Fig. 4B). We notice there is a set of causal edges shared by tumors in Cluster 1 (red box in Fig. 4B and 4C) that is absent in other Clusters, including Cluster 2, which is enriched with CMS4 samples. Edges in this block are enriched with interactions involving CAFs, endothelial cells, and immune cells, indicating divergent underlying communication programs. This suggests that iGFCI not only unveils the unique communication patterns distinguishing CMS4 subtype from CMS1-3 subtypes, but also provides information complementary to traditional CMS subtypes. It can uncover meaningful subgroup heterogeneity within the poor-outcome CMS4 subtype, with potential relevance for personalized therapeutic targeting.

### 3.3. Precision stratification of TCGA CRC CMS4 patients using tumor-specific CCCNs

While scRNA-seq datasets in this study provide high-resolution insight into cellular states and interactions, they lack long-term clinical outcomes such as overall survival. To evaluate the clinical relevance of GEM-GEM causal edges identified by iGFCI, we leveraged the TCGA CRC bulk RNA-seq dataset with corresponding clinical outcome data. The TCGA cohort offers an opportunity to validate the identified cell-cell communication patterns and assess their prognostic significance.

#### 3.3.1. Stratification based on causal edges and their associated GEMs in TCGA CRC cohort

Besides representing a tumor as a binary vector (Fig. 4B), indicating the presence or absence of a causal edge in a tumor, we further integrate the expression status of GEMs connected by a causal edge to provide richer information related to an edge. This is based on the hypothesis that the expression status of a GEM reflects the state of the signaling pathway that regulates its expression in a subpopulation of cells. For each GEM, we represented its expression status by taking the average expression of the top 50 genes associated with it (Supplementary Table 1). Next, for each causal GEM pair, we computed the geometric mean of the two corresponding GEM expression values. We called the calculated geometric means causal edge scores (CES). These values were then z-scored across all patients to create a normalized matrix representing the activity of each causal edge.

We calculated the CESs of the causal edges highlighted in the red box (edges of interest, EOIs) of Fig. 4C for all TCGA tumors. Using this as representation, we performed clustering which divided patients into four groups (Fig. 5A). The survival is significantly different among these four clusters (p < 0.05; Fig. 5B). Cluster 1, characterized by high activity of EOIs, exhibited significantly worse overall survival compared to other clusters (p = 0.01; Fig. 5C). These results demonstrate that intercellular signaling patterns inferred using iGFCI may serve as informative prognostic markers for stratifying CRC patients.

**Fig. 5.**
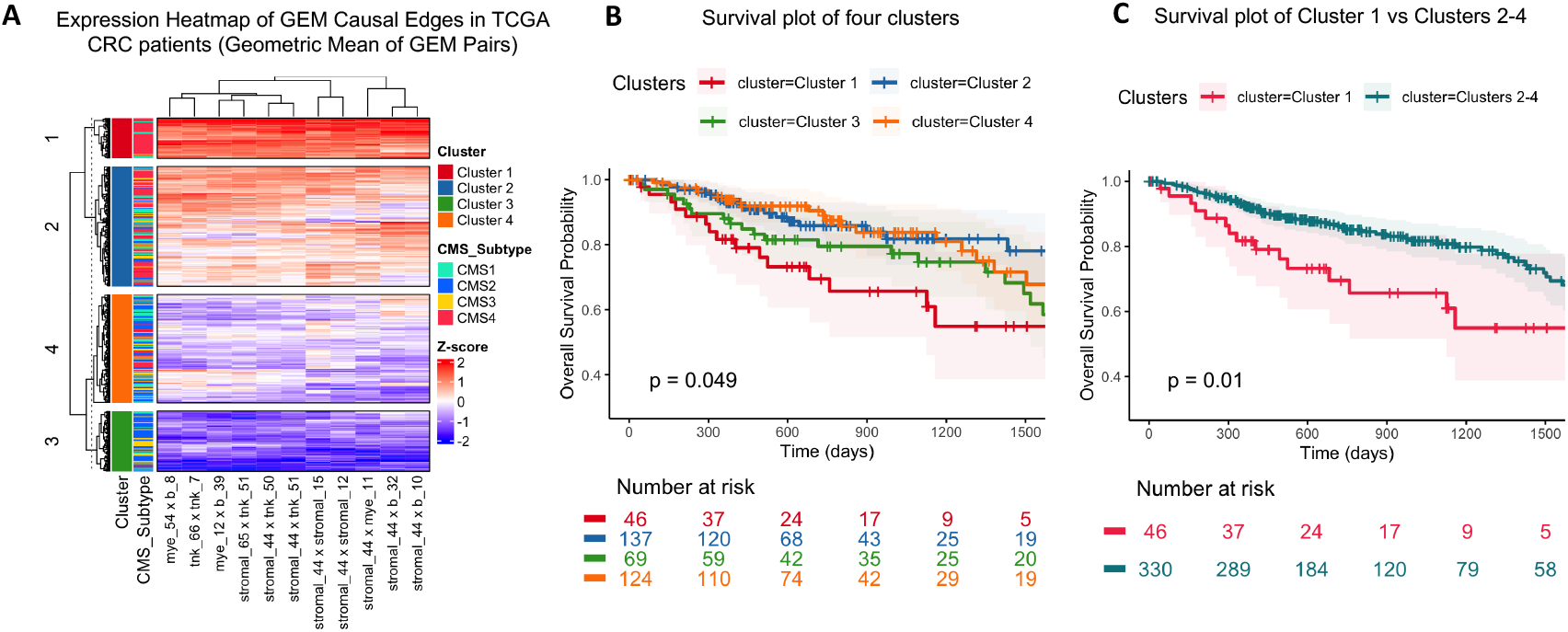
Stratification of CRC patients in TCGA using GEM causal edges. **A**. Heatmap of GEM causal edge expression across all TCGA CRC patients. The expression values are determined by the z-scored CES of GEM pairs involved in EOIs (from Fig. 4C). Patients are annotated by K-means clustering (K=4) and CMS subtypes. **B**. Kaplan-Meier survival curve comparing four clusters in TCGA CRC patients (p < 0.05). **C**. Kaplan-Meier survival curve comparing Cluster 1 with other clusters in TCGA CRC patients (p = 0.01).

#### 3.3.2. Uncover CMS4 heterogeneity via causal edges

Studies have shown that CRC patients in the CMS4 subtype have worse outcomes compared to other CMS subtypes^35^, potentially due to poor anti-tumor immune response. We focused on TCGA CMS4 patients (n=126) to examine whether distinct TMEs, reflected by individualized CCCNs, can shed light on the disease mechanisms of patients in this subgroup. We calculated the CES of EOIs and performed clustering analysis, which yielded four patient clusters (Fig. 6A), with Cluster 2 exhibiting high CES expression of EOIs.

**Fig. 6.**
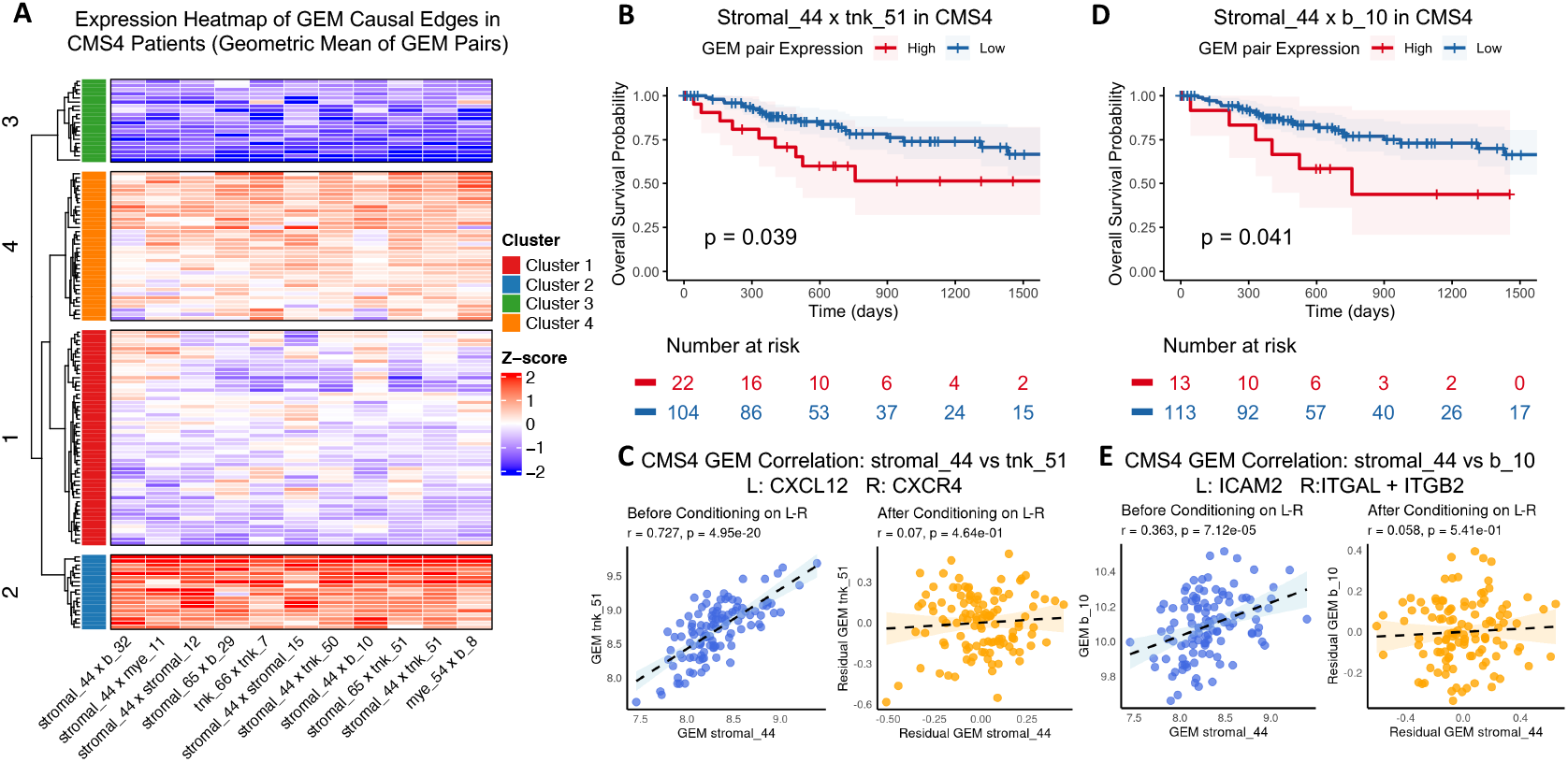
Prognostic value and biological validation of specific GEM causal edges in TCGA CMS4 patients. **A**. Heatmap of GEM causal edge expression (z-scored CES of GEM pairs enriched in CMS4 group 2 as shown in Fig. 4C) in CMS4 patients. Patients are grouped into four clusters. **B**. Kaplan-Meier survival curve for CMS4 patients stratified by stromal_44 × tnk_51 expression. High expression is associated with poor prognosis (p = 0.039). **C**. Scatter plots showing GEM correlation between stromal_44 and tnk_51 (left), and residuals correlation after conditioning on the LR pair CXCL12-CXCR4 (right). **D**. Kaplan-Meier survival curve for stromal_44 × b_10 expression stratification. High expression again predicts worse survival (p = 0.041). **E**. Correlation and conditional independence analysis for stromal_44 and b_10 conditioned on LR pair ICAM2–ITGAL+ITGB2.

We also examined the association of the CES of an individual causal edge and patient outcomes. The stromal_44 × tnk_51 interaction, representing endothelial to CD4^+^ Th1 cell communication, was associated with worse prognosis (p = 0.039; Fig. 6B). Similarly, stromal_44 × b_10, reflecting endothelial to IgA B cell communication, was also predictive of poor outcome (p = 0.041; Fig. 6D).

### 3.4. Investigating molecular mechanisms of CCC captured by causal edges

A causal relationship between two types of cells expressing corresponding GEMs in one GEM-GEM causal edge indicates that the two cell types communicate with each other, which involves LR interactions to transmit signals from one cell to another. To search for LRs that potentially mediate signal transduction between cells expressing the GEMs connected by a causal edge, we performed conditional independence testing using LR pairs from the CellPhoneDB v5 database (Supplementary Table 2). As an example, we found that the correlation between stromal_44 and tnk_51 GEMs (r = 0.727) was substantially reduced after conditioning on the CXCL12-CXCR4 pair (Fig. 6C), suggesting this LR interaction potentially mediates the observed causal edge.

Biologically, stromal_44 is expressed predominantly in endothelial cells and reflects a program of angiogenesis, EndoMT, and metabolic stress, marked by genes such as *DLL4*^25^, *SNAI1*^26^, and *POLD4*^28^. It is highly correlated with tnk_51, which suggests that the cells expressing stromal_44 interact with the cells expressing tnk_51. Tnk_51 is a GEM enriched in CD4^+^ Th1 cells expressing immune-related genes (*CD40LG*^36^, *TNFSF13B*^*37*^, *FLT3LG*^38^), which may reflect endothelial-driven recruitment and activation of effector T cells within the tumor microenvironment. The CXCL12-CXCR4 axis, involving CXCL12 produced by endothelial cells and CXCR4 expressed on immune cells, is a well-established chemoattractant pathway that facilitates T cell trafficking into peripheral tissues, including tumors^39^. However, in the CMS4 subtype of colorectal cancer, characterized by a dense stroma, active angiogenesis, and immunosuppression, activation of this signaling axis may trap T cells in perivascular or stromal compartments, limiting their access to tumor cells, and potentially promoting T cell dysfunction or exhaustion, thereby contributing to poor prognosis^40^. Studies have shown that *DLL4*^41^, *SNAI1*^42^, and *POLD4*^43^ from stromal_44, the ligand-receptor pair CXCL12–CXCR4^44,45^, and *CD40LG*^36^, *TNFSF13B*^46^, and *FLT3LG*^47^ from tnk_51 are all individually associated with poor prognosis in cancer patients and have the potential to be prognostic biomarkers. The GEM causal edge between stromal_44 and tnk_51 identified in our study may provide a mechanistic explanation for how these prognostic biomarkers interact and contribute to adverse clinical outcomes.

Similarly, the stromal_44 and b_10 interaction, reflecting endothelial to B cell communication, also showed a significant correlation (r = 0.383), which was largely removed after conditioning on the ICAM2–ITGAL+ITGB2 LR complex (Fig. 6E). This implies that this integrin-mediated adhesion pathway may underlie the causal relationship. Biologically, b_10 is a GEM enriched in IgA^+^ B cells, associated with oxidative metabolism and protein secretion, involving genes like NDUFV2^48^, IDH2^49^, and SSR1^50^. These transcriptional features align with plasma-like B cells that support immunoglobulin production and potentially enhance tumor-promoting inflammation or immune suppression. ICAM2–LFA-1(ITGAL+ITGB2) interactions promote immune cell adhesion and positioning within tissue compartments^51^. In CMS4 tumors, the coupling of endothelial remodeling (via stromal_44) with immunosuppressive IgA^+^ B cell phenotypes (via b_10) may contribute to sustained immune evasion and tumor progression, consistent with worse survival observed in patients with high expression of this GEM pair.

## 4. Discussion

In this study, we present a computational framework that overcomes key limitations of existing CCC analysis methods by constructing personalized, causal communication networks for each patient. Traditional CCC tools often rely on population-level correlations and cluster-averaged LR signals, which can obscure individual variability. In contrast, we use iGFCI to infer directed causal relationships between GEMs in each tumor, moving beyond correlation-based links to reveal mechanistic interactions. Our method leveraged hierarchically structured GEMs via the nested hierarchical Dirichlet process, and our approach achieved a high-resolution view of each tumor’s microenvironment. By integrating large-scale scRNA-seq data (so far the largest collection of CRC scRNA-seq data) at detailed cell subtype resolution, this individualized strategy yields greater precision for CCCN inference and allows us to stratify patients into novel subgroups that extend beyond traditional classifications like the CMS subtyping.

Our framework introduces several innovations. First, it provides a new causality perspective to investigate CCCNs, particularly individualized CCCNs. This framework provides more mechanistic insights into CCCNs that define TME, in comparison to conventional clustering-based methods of studying CCC. Each patient’s CCCN is inferred separately, preserving heterogeneity that would be lost by pooling population-wide data to perform analysis. Second, using a causal inference approach provides directed edges that imply upstream-downstream influences, enabling mechanistic insights beyond what correlation-based methods offer. We provide evidence that some of the discovered causal edges are further supported by LR signaling (e.g., conditioning on CXCL12–CXCR4 interaction explains the correlation between stromal_44 in endothelial and tnk_51 in CD4^+^ Th1 cells; Fig. 6C). Third, our method provides a new perspective to stratify patients based on CCCNs. Our method leads to novel stratification of CRC patients, which revealed a potential immune evasion mechanism that could not be detected by CMS classification. The clinical utility of this novel perspective remains to be further investigated. The impact of this approach is highlighted in CMS4 CRC, a subtype associated with dense stroma, immunosuppression, limited response to available therapies including immunotherapy, and poor prognosis^35^. Our causal discovery results revealed the immune evasion mechanisms that can be targeted by new medicine.

Our method could be further improved. Stable causal inference typically requires large sample sizes (cells per tumor and patient cohorts) to ensure robust identification of interactions. And further testing on a larger dataset may be required to produce more conclusive results. Future work will include integrating spatial transcriptomics to investigate cell–cell interactions in their tissue contexts, confirming physical colocalization of interacting cells. Additionally, applying additional causal discovery algorithms into our framework, such as BOSS Fast Causal Inference (BFCI)^52^, could potentially enhance network accuracy and complexity modeling. Extending this framework to additional cancer types and incorporating other data modalities (e.g., proteomics, epigenomics) will also help generalize and deepen understanding of intercellular communication networks. In conclusion, our individualized causal inference approach provides valuable insights into tumor heterogeneity and highlights a new way toward precision oncology.

## 5. Acknowledgments

This research was supported in part by the National Library of Medicine, National Human Genome Research Institute, National Cancer Institute at the National Institutes of Health grants R01LM012011, R00LM013089, R01HG014023, and R01CA254274.

## 6. Supplemental Materials

All supplemental materials are available at: https://github.com/pikapika-qiu/CRC_causal_iCCCN.git

